# SmartGraph: A Network Pharmacology Investigation Platform

**DOI:** 10.1101/707869

**Authors:** Gergely Zahoránszky-Kőhalmi, Timothy Sheils, Tudor I. Oprea

**Affiliations:** National Center for Advancing Translational Sciences, Rockville, MD; Department of Internal Medicine, University of New Mexico School of Medicine, Albuquerque, NM, USA; UNM Comprehensive Cancer Center, Albuquerque, NM, USA; Department of Rheumatology and Inflammation Research, Institute of Medicine, Sahlgrenska Academy at University of Gothenburg, Gothenburg, Sweden; Novo Nordisk Foundation Center for Protein Research, Faculty of Health and Medical Sciences, University of Copenhagen, Copenhagen, Denmark

## Abstract

**Motivation:** Drug discovery investigations need to incorporate network pharmacology concepts while navigating the complex landscape of drug-target and target-target interactions. This task requires solutions that integrate high-quality biomedical data, combined with analytic and predictive workflows as well as efficient visualization. SmartGraph is an innovative platform that utilizes state-of-the-art technologies such as a Neo4j graph-database, Angular web framework, RxJS asynchronous event library and D3 visualization to accomplish these goals.

**Results:** The SmartGraph framework integrates high quality bioactivity data and biological pathway information resulting in a knowledgebase comprised of 420,526 unique compound-target interactions defined between 271,098 unique compounds and 2,018 targets. SmartGraph then performs bioactivity predictions based on the 63,783 Bemis-Murcko scaffolds extracted from these compounds. Through several use-cases, we illustrate the use of SmartGraph to generate hypotheses for elucidating mechanism-of-action, drug-repurposing and off-target prediction.

**Availability:** https://smartgraph.ncats.io/

## Introduction

The path to successful drug discovery frequently demands understanding the complex network-pharmacology (NP) landscape [1]. Understanding how the action of a drug perturbs the intricate regulatory relations among protein targets is pivotal to surmounting this challenge. Thus, it is imperative to associate drugs, targets, clinical outcomes and side effects within an integrated network [2]. Specifically, for drug-target interactions (DTIs), NP opens the possibility of evaluating potential DTIs in the context of their complex biological and chemical setting. Adopting and utilizing NP-based workflows in everyday research remains a challenging task, yet one that is likely to lead to more fruitful scenarios.

A rational design workflow that integrates NP needs to consider multiple interactions between a drug molecule and its intended (target, or “on-target”) and not-intended (“off-target”) interaction partners. Here, we consider proteins to be the key DTI partners. Due to the existence of multiple pathways between proteins, the effects of a drug molecule might ripple to proteins not in direct contact with the drug in question. Such perturbations may also alter gene-expression profiles of cells. The complex nature of this interaction-landscape demands novel and integrated target-chemistry-bioactivity approaches [3] in drug discovery and facilitates the devising of new therapeutic strategies, e.g., multitarget therapies. Therefore, it is imperative to develop new ways of analyzing such complex relationships. The ever-growing nature of biomedical data demands the use of modern technologies such as graph-databases, to facilitate the analysis, processing and retrieval of drug-target and target-target relations at scale. A computational NP framework addressing these aspects is essential in the quest to translate biomedical information into clinically relevant knowledge. Here we present such a framework: *“SmartGraph”*, which has two aims:

First, create an easy-to-use graphical user interface (GUI) that can facilitate various steps of the drug discovery process without requiring in-depth expertise in bioinformatics and cheminformatics. Target deconvolution of phenotypic assays, on- and off-target identification, lead and chemical probe discovery as well as drug repurposing are among the major activities that SmartGraph can support. Second, SmartGraph facilitates the development of predictive and analytic NP workflows.

Other network-analysis inspired tools can facilitate the analysis of DTIs and PPIs, such as the CARLSBAD Web Application and Cytoscape plugin [4], the ChemProt database [5], the STRING-based STITCH platform [6], [7], the Drug Target Explorer [8], Bioactivity-explorer [9], and the PathInsight [10] and PathLinker Cytoscape plugins [10], [11]. SmartGraph combines some of the distinctive characteristics of these platforms: It integrates the use of common chemical patterns as implemented in CARLSBAD and CarlsbadOne [12], the pathway analytic features of STITCH and STRING, the ability to define starting and end nodes (PathLinker), the integration of DTIs with PPIs (PathInsight) and the ability to predict new drug-target interactions (CarlsbadOne and Drug Target Explorer). Bioactivity-explorer has some similar functionalities with SmartGraph, mainly the use of scaffolds for relating targets but it lacks the integration of pathway analysis and bioactivity prediction. In addition, SmartGraph is built on Neo4j [13], a state-of-the-art graph database, and uses their bolt websocket interface [14] for fast data transfer. Angular is used to generate object models and UI templates, and RxJS [15] is used to asynchronously update the data retrieved from Neo4j. This data is coupled with a robust D3.js force-directed graph visualization to efficiently handle and display this large-scale data, and to support the integration of additional layers of biomedical data [16][17]. Finally, distinctive features of SmartGraph compared to existing tools are a Google-Maps-like [18] “start and destination”-based pathway discovery feature, and an integrated bioactivity prediction algorithm based on target-class agnostic “potent chemical patterns” [19]. The use of the Neo4j database engine enables these features to work on a large scale of data and in a practically instantaneous manner, unlike existing tools.

## Computational Methods and Datasets

### Rationale

With compounds and targets as nodes and relationships as edges, we can model DTIs and PPIs as a network. An edge linking a compound and a target could represent an experimentally determined bioactivity value, whereas an edge between two targets could be a regulatory relation between the two. Nodes and edges can have various attributes, such as the IC_50_ (bioactivity) value or the specific regulatory mechanism linking two targets. From a structure-activity-relations (SAR) standpoint, targets tend to recognize certain chemotypes, i.e. common chemical patterns in a series of molecules. Derived from known active compounds, chemotypes can be exploited in the prediction of novel DTIs. The Bemis-Murcko scaffolds [20] are examples of chemotypes, but other algorithms could be used for this task, such as Maximal Common Edge Subgraph algorithm [21], Scaffold Network Generator [22] and Hierarchical Scaffolds [23], [24] to name a few. We refer to chemotypes as ‘patterns’ throughout this text to imply the general nature of these objects. Detailed descriptions of object (node) and relation (edge) types are provided in the “Terminology” section of SI (Supplementary Information).

### Preprocessing DTI and PPI Information

Information for compounds, targets and their interactions were extracted from the ChEMBL (version: 24.1, downloaded on: 10/26/2018) [25], [26] and SIGNOR (version: 2.0, Homo Sapiens, downloaded on: 04/23/2017) [27] databases. With the help of a KNIME [28] workflow (detailed in SI) bioactivity data related to human targets were aggregated by unique compound-target-bioactivity type triples, in conjunction with Bemis-Murcko pattern detection (see “Bioactivity Data Aggregation” SI section). The aggregated bioactivity values represent the median value of all observed bioactivities for a given triple.

### Data Integration

Pre-processed DTI and PPI information (compounds, targets, patterns and their relationships) was then assembled into a backend Neo4j database, via JDBC-based Java code connecting the PostgreSQL [29], [30] (ChEMBL and SIGNOR) and Neo4j [13], [31] databases; the code is available publicly at https://github.com/ncats/smartgraph_backend.git upon the acceptance of this manuscript. This repository contains an SQL script to create a database and load corresponding data. As input, the script uses KNIME-created files [28]. The Java code then uses DTI compounds, and DTI and PPI targets, represented by their InChI [32] keys and UniProt IDs [33], respectively. For convenience, the non-stereo part of the InChI key (NS-InChI key) of compounds are also provided as a compound-attribute, see: “Terminology” (SI). Each node and edge in the Neo4j database is assigned a unique UUID identifier [34] for efficient data handling and database queries.

### Metadata

SmartGraph includes metadata associated with compounds, patterns, targets and their relationships as node and edge attributes, respectively. Such attributes include the median activity between a compound and a target (when multiple activities are available), the confidence value of a given PPI, its nature (e.g., phosphorylation, ubiquitination, etc.), and the overlap of a pattern with the associated compound as the ratio of their heavy atoms. Compound and pattern nodes are also annotated with the respective SMILES and structure depictions. Target nodes include gene symbols and names as attributes as well.

### Network Perturbation

To interpret phenotypic screening results, one can use SmartGraph to find all shortest paths between user-defined start and end nodes. When multiple such shortest paths are found, all connecting shortest paths between the selected two nodes are returned. Once merged into sub-networks, these shortest paths can be used to generate hypotheses for the explanation of mechanism-of-action of phenotypic screen hits. Despite the large amount of information stored in the backend database, this operation is performed efficiently by the underlying Neo4j database engine using the Cypher query language [13] and bolt websocket interface [14].

When the activity of a protein is perturbed, downstream proteins within the same pathway are affected as well. Given the complex nature of each pathways, such perturbations affect targets at increasing distances in a less and less deterministic manner. To account for this phenomenon, the parameter (*k*) introduced into the shortest path detection process limits the length of shortest paths in connecting start and end nodes. By default, *k* = 5.

Start nodes can be defined in two ways: First, a set of small molecules with their associated NS-InChI keys can be start nodes. In such cases, SmartGraph identifies proteins on which any of these small molecules were assayed using the DTIs extracted above. An additional parameter (*p*) can be used to filter DTIs that are of potency lower than *p* (in negative log units). Hence, proteins associated with DTIs of AC_50_/IC_50_/EC_50_ below *p* will not be considered as start nodes after filtering. By default, *p* = 5. Second, UniProtID IDs for proteins can also be used. SmartGraph supports mixed queries, i.e. input sets can contain both small molecules and proteins as start nodes.

The shortest path exploration process can be influenced by an additional parameter (*c*), regardless of the start node definition method. The SIGNOR database stores confidence (*c*) values for each PPI. By adjusting the value of *c*, low confidence PPIs can also be excluded from the shortest path discovery process.

It is possible to leave the end nodes undefined, which enables manual exploration of pathway perturbations. SmartGraph includes features to expand the node neighbors list on a node-type basis. For instance, in the case of a protein node it is possible to expand compound/protein/pattern neighbors. It is also possible to selectively hide neighbors based on node types.

### Bioactivity Prediction

Experimental bioactivities are often used to classify compounds as active or inactive. The cut-off value for such categorization is often context-dependent, particularly with respect to target type, and the wealth of information (e.g., number of compounds with measured bioactivities) available at any given time. The active/inactive separation may further depend on SAR, as some chemotypes may be considered more active than others.

In seeking to emulate a medicinal chemist’s approach, we decided to classify the top 20% of the compounds (ranked in decreasing order of bioactivity) as “potent”, with the remaining compounds tested on that same target being labeled as “not potent”. This approach is preferable when working with multiple target classes, since the typical enzyme or ion channel bioactivity profile may vary by two or three orders of magnitude compared to a nuclear or G-protein coupled receptor. Thus, the 0.1*μ*M cut-off value (7 on the –log scale) does not always appropriately separate actives from inactives across all target classes. The 80/20 principle was generally applied to partition compounds, but for certain proteins, an alternative formula was required. When a protein had at most five (aggregated) bioactivity values, the cutoff value was set to 7 (0.1*μ*M). Cutoff values were lumped across all bioactivity values, regardless of type, e.g. AC_50_, IC_50_, EC_50_. Although some might use K_i_ as surrogate for IC_50_/AC_50_/EC_50_ values, in this study we did not consider K_i_ or other type of bioactivity values besides IC_50_/AC_50_/EC_50_ types.

Potent compounds for each target were tagged using bioactivity values from ChEMBL and the above criteria. We then examined the relationship between proteins based on their molecular recognition ability, estimated by chemical patterns (here: scaffolds) featured in potent compounds. These target-compound and compound-pattern associations allow us to infer novel associations between targets and patterns. All patterns contained in a potent compound can thus be considered *potent patterns* for that specific target. Associations between a target and potent patterns suggest that certain targets might be specifically perturbed by other (untested) compounds featuring the same potent patterns.

## Results and Discussion

### Knowledge Base of SmartGraph

The knowledgebase of the framework was implemented in Neo4j graph-database which contains 1,732,553 unique compounds and 4,583 protein targets. With the help of Bemis-Murcko scaffold analysis, we extracted 523,944 unique scaffolds from compounds. We observed that some of these scaffolds are connected to too few or too many compounds. In order make the bioactivity prediction efficient, we decided to remove scaffolds deemed too general or too specific from the knowledge base, i.e. scaffolds connected at least to 100 or to less than 5 compounds, respectively. This process gave rise to 63,783 unique scaffolds, i.e. patterns, that participate in 793,085 one-to-many pattern-compound relations. Furthermore, 271,098 out of all compounds participate in 420,526 DTIs involving 2,018 protein targets. Out of these targets, 1,455 are associated with a potent pattern and 901 participate in PPIs, see: *Figure 1*. All information about compounds, targets and scaffolds is stored in the Neo4j graph-database.

**Figure 1.**
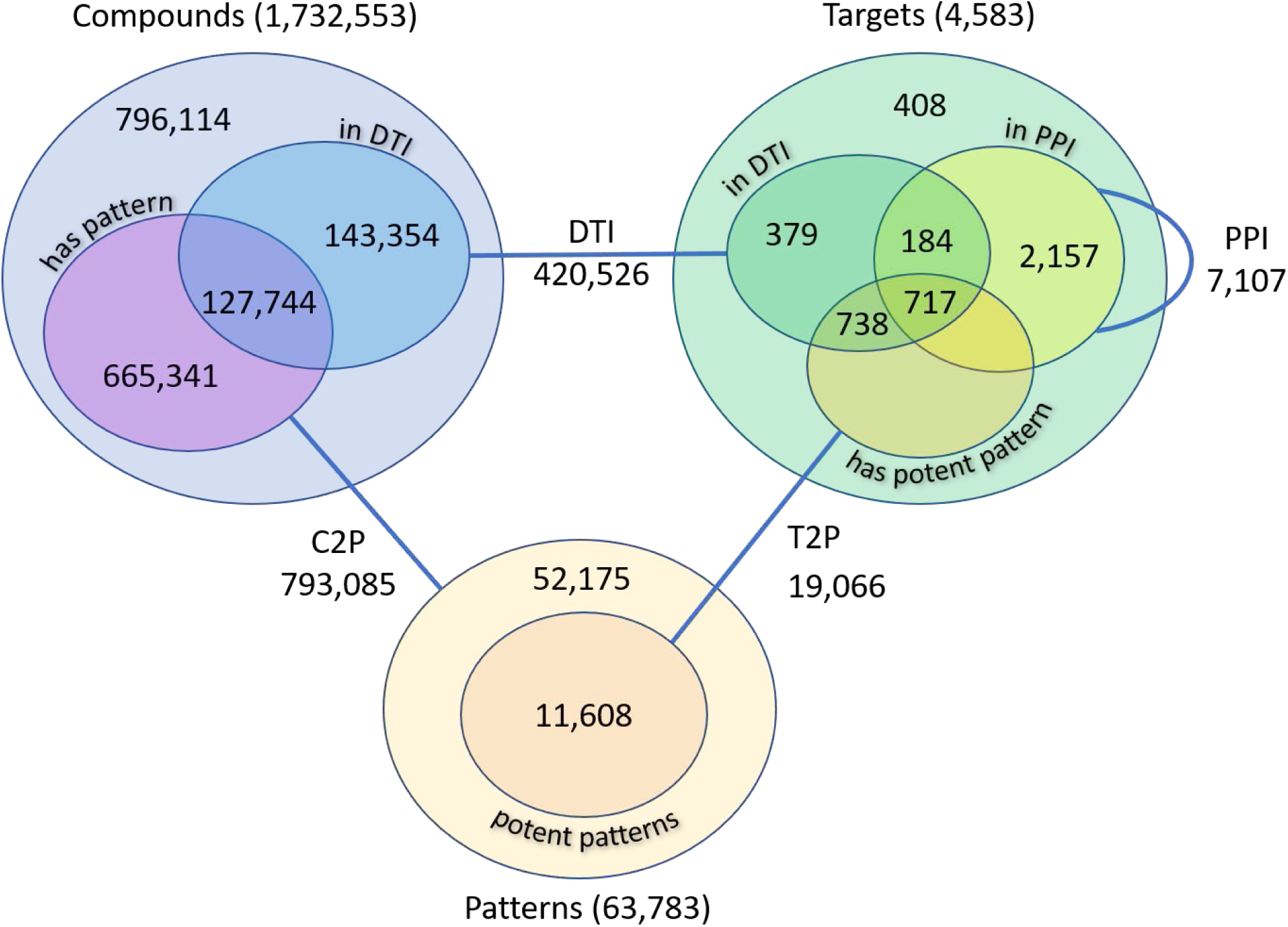
Knowledge base of SmartGraph. C2P: pattern-of relations, T2P: potent pattern-of relations, DTI: drug-target interactions, PPI: protein-protein interactions.

### Graphical User Interface of SmartGraph

The network analysis and bioactivity prediction methods of SmartGraph are exposed through a web-based GUI available at https://smartgraph.ncats.io/. We recommend Chrome, Firefox and Safari browsers to use SmartGraph. This GUI was implemented using the D3 visualization library to generate a force directed graph interface, and the Angular [35] framework to track and update the graph data as it changed. The GUI also uses RxJS, an asynchronous event programming library to set up a websocket to communicate with the backend server through the Neo4j “bolt” protocol [14], making the queries much faster than if they were done through traditional API calls. The GUI itself is organized into three main panels (see: Figure 2).

**Figure 2.**
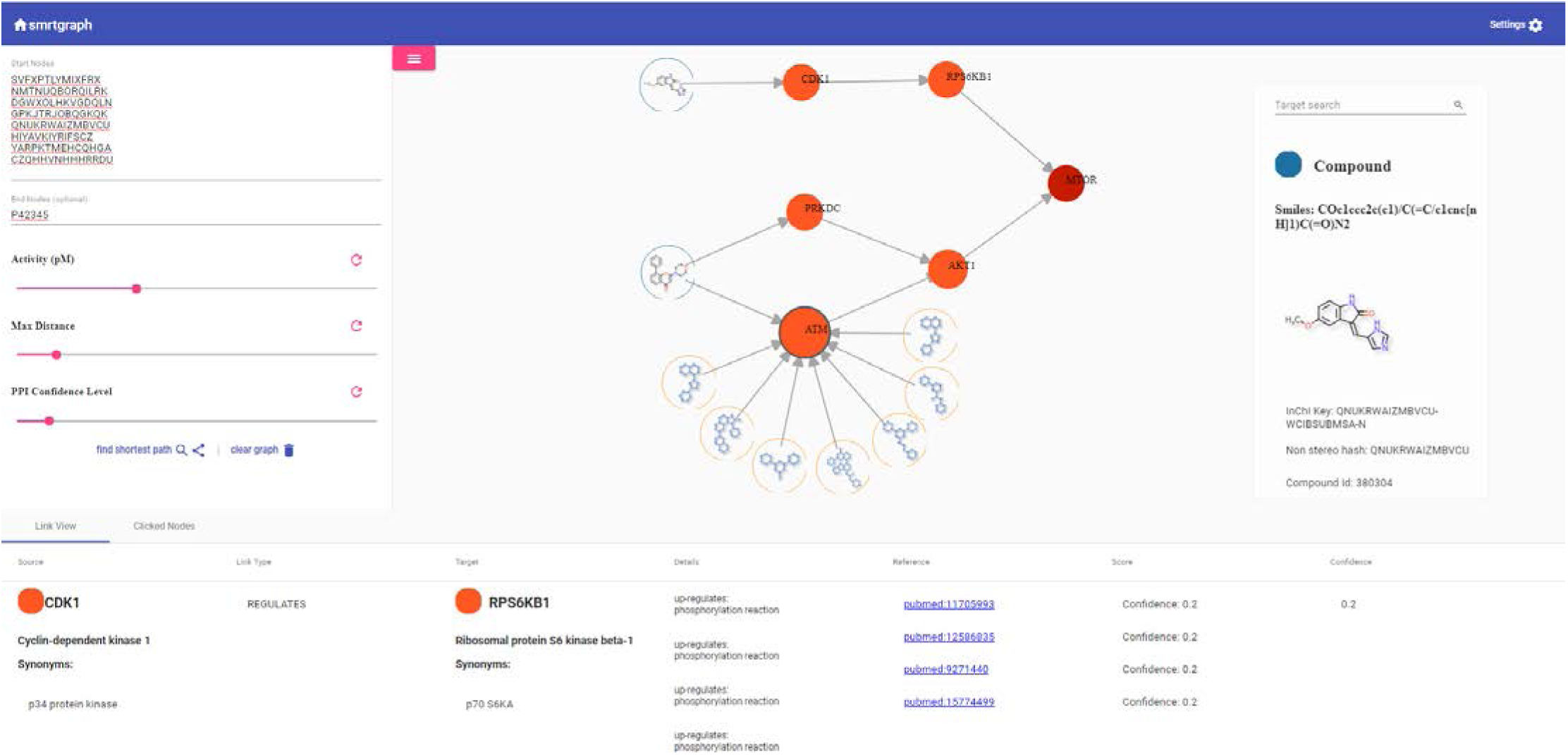
Web-based Graphical User Interface (GUI) of SmartGraph. Shown is an example network that was generated by 8 compounds and one destination target and after adjusting parameters *k=2*, *p=1uM* and *c=0.10* in the control panel. The structures of two compounds are oriented on the left-hand side of the subnetwork. The structures of pattern nodes (BM-scaffolds) are outlined in blue, and their nodes are embordered by a yellow line. The end node is colored by dark red, whereas other targets are colored by orange. These targets constitute the shortest paths between the compounds and the destination target.

The control panel (left side) provides the means to construct and fine-tune subnetworks for pathway analyses and is collapsible to allow for a larger view of the generated network. The selection of start and end nodes and the values of *k*, *p* and *c* can be set using this panel. The main (visualization) panel is used to visualize and interact with the generated subnetworks, clicking on a node reveals a floating menu with the following features: Expand/collapse compound/pattern/target/all neighbor nodes; and generate bioactivity predictions, respectively. The main panel also contains a smaller sub panel, which shows node details on hover, and allows the user to search for protein nodes within the displayed graph and isolate the nearest neighbors. The third, bottom (information) panel is used to visualize selected node and edge metadata in two separate tabs. Information is displayed in this table upon hovering over edges or by clicking on nodes. Clicking on a node or edge will persist it in this table, allowing users to generate a list of interesting nodes or edges. Some of the information displayed is the depiction of chemical structures and patterns, bioactivity in μM unit, and confidence of PPIs, to name a few.

The information displayed on the nodes can be configured via a collapsed menu accessed by clicking on the gear icon on the top-right corner of the visualization panel. The user can set the label type on various nodes: protein/gene name, symbol, identifier and compound/pattern identifier or structure. This menu also contains a legend denoting the different node types as well. The control panel can be collapsed by clicking on the icon with three bars on the top-left corner of the visualization panel to provide more visual real estate for the graph visualization. Finally, an additional panel overlaid on the network-panel provides means to search for target nodes in the network based on their names and to lookup their record in the Pharos database [36] via link-out. The same panel also provides the most important information of compound and pattern nodes. The source code used to generate the GUI is available publicly at https://github.com/ncats/smartgraph_frontend upon the acceptance of this manuscript.

### Use Cases

To illustrate some of the features and typical use cases of the SmartGraph platform, we selected an mTOR signaling pathway assay (PubChem [37], [38] AID: 2660), using the qHTS technology [39] in confirmatory mode, i.e., dose-response using 7 concentration points. mTOR is linked to a number of diseases, including cancer [40]. The assay identified a downstream effect of mTOR activation. Out of 1,280 compounds, 8 were confirmed actives. Below, we describe three typical use-cases for SmartGraph.

### Manual Exploration of Biological Relations

When active compounds are identified via a phenotypic screening, one of the next steps is to investigate targets associated with these actives and to try to link them to other targets that may play important role in the observed phenotype. Such an exploratory analysis can be initiated by providing the NS-InChI key of active compounds in the “Start Nodes” field of the control panel or by inputting the UniProt identifier of known targets. Here we use a selected active compound (NS-InChI key: DGWXOLHKVGDQLN, PubChem CID: 398148) as a starting point. The identified start node(s) appear in the visualization panel (compound and target nodes shaded blue and orange, respectively). Next, when left-clicking on the nodes a floating menu with several options appears. Manual exploration of relevant biological relationships can then be performed using the “Expand Target” feature, which in this case reveals three targets for which the compound at hand was tested: CDK1, CDK2 and ATR (gene symbols). On the information panel the bioactivity value of a DTI can be revealed by clicking on the “Link View” tab and hovering the mouse over, or clicking on the edge defined between the respective compound and the target. The result of further expanding this network on ATR target is shown in *Figure 3*.

**Figure 3.**
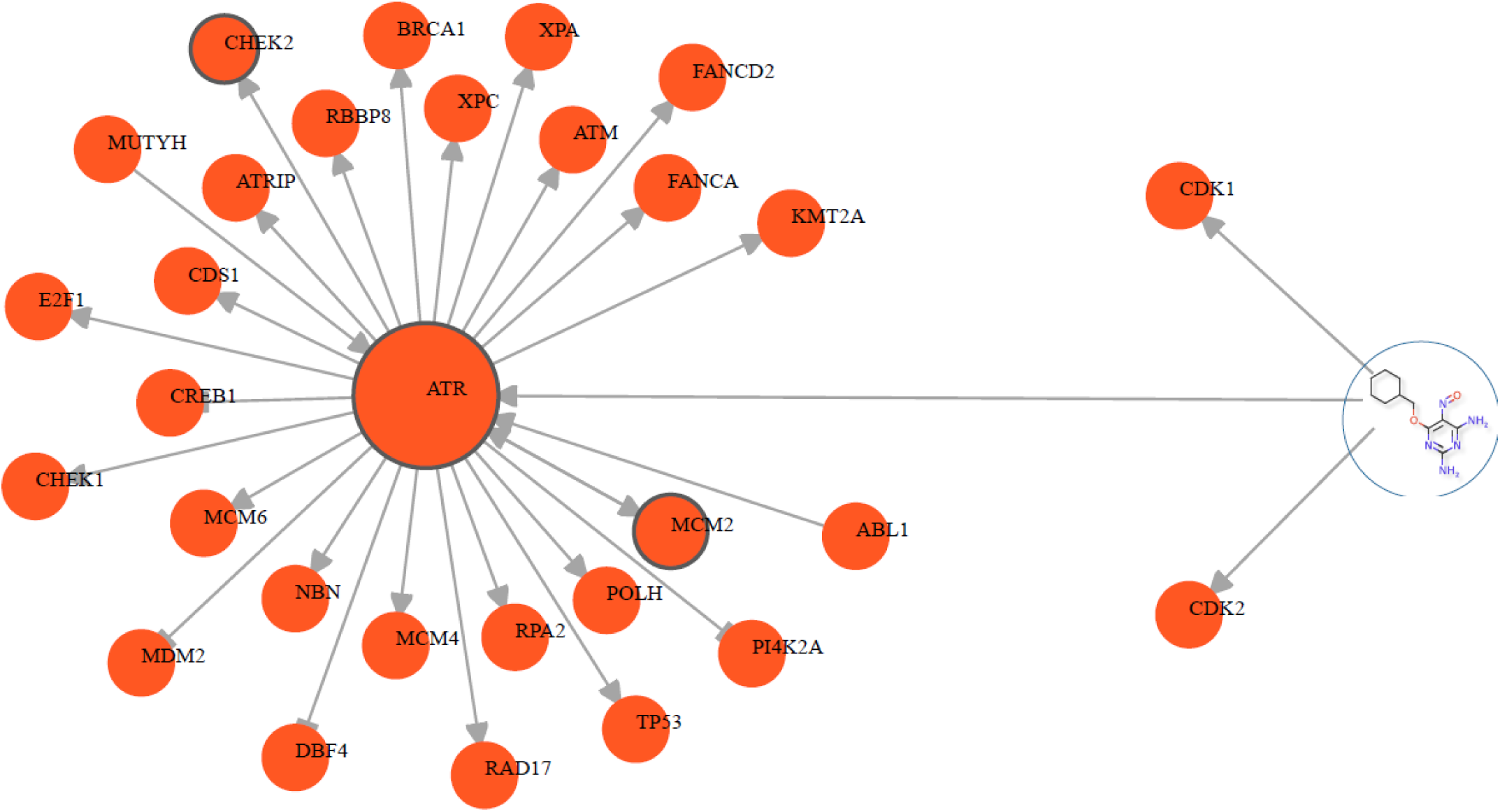
First use-case. Shown is a subnetwork that can be revealed by tracking the effect of a compound by the means of manual exploration. The network was assembled by defining a compound as a starting node (PubChem CID: 398148), and subsequently the neighboring target nodes of the compounds were expanded. Finally, the neighboring target nodes of ATR protein target, as a potential target of interest, were also expanded.

### Automated Hypothesis Generation for Mechanism-of-Action

While the first use-case can be performed using multiple start nodes, the process can be laborious given that some of targets are involved in multiple PPIs. As described in the “Network Perturbation” section, SmartGraph can reveal shortest path(s) between a start node and an end node. Such an analysis can be initiated, for example, by entering a space/new-line/comma-separated list of the NS-InChI keys of the 8 active compounds pasted into “Start Nodes” and the UniProt identifier of mTOR (P42345) into “End Nodes”, given that the assay at hand interrogates mTOR activation. We recommend clicking the “clear graph” button, followed by re-inserting the respective node identifiers when one wants to re-define start or end nodes. After clicking the “find shortest path” button, SmartGraph will attempt to find shortest paths between the two sets using the default values of parameters. By setting the three filtering parameters to permissive values, i.e. *k=5*, *p=3*, *c=0.01*, the resulting subnetwork includes three compounds with multiple shortest paths. This subnetwork and subsequent ones in this section are shown in *Figure 4*.

**Figure 4.**
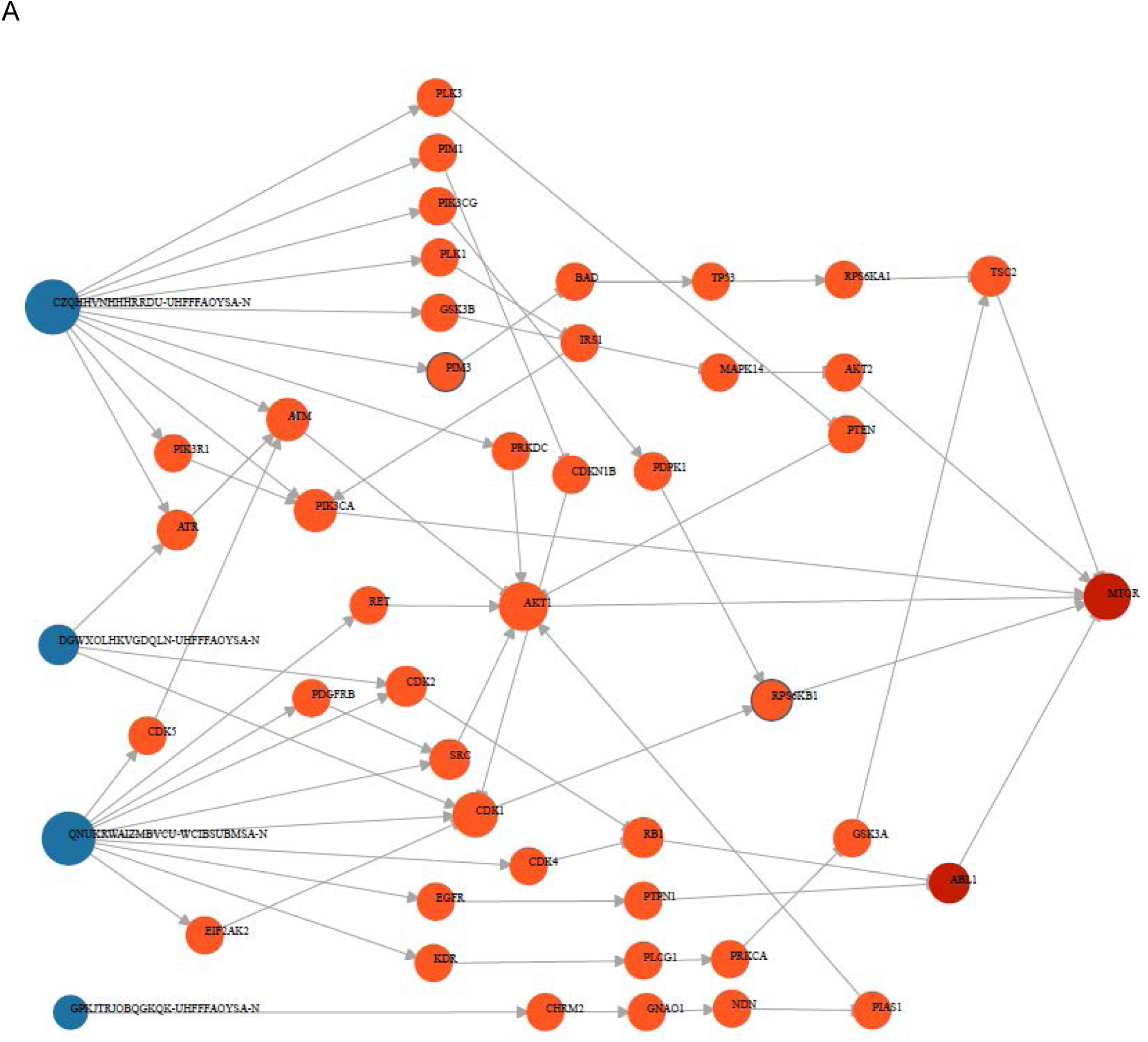
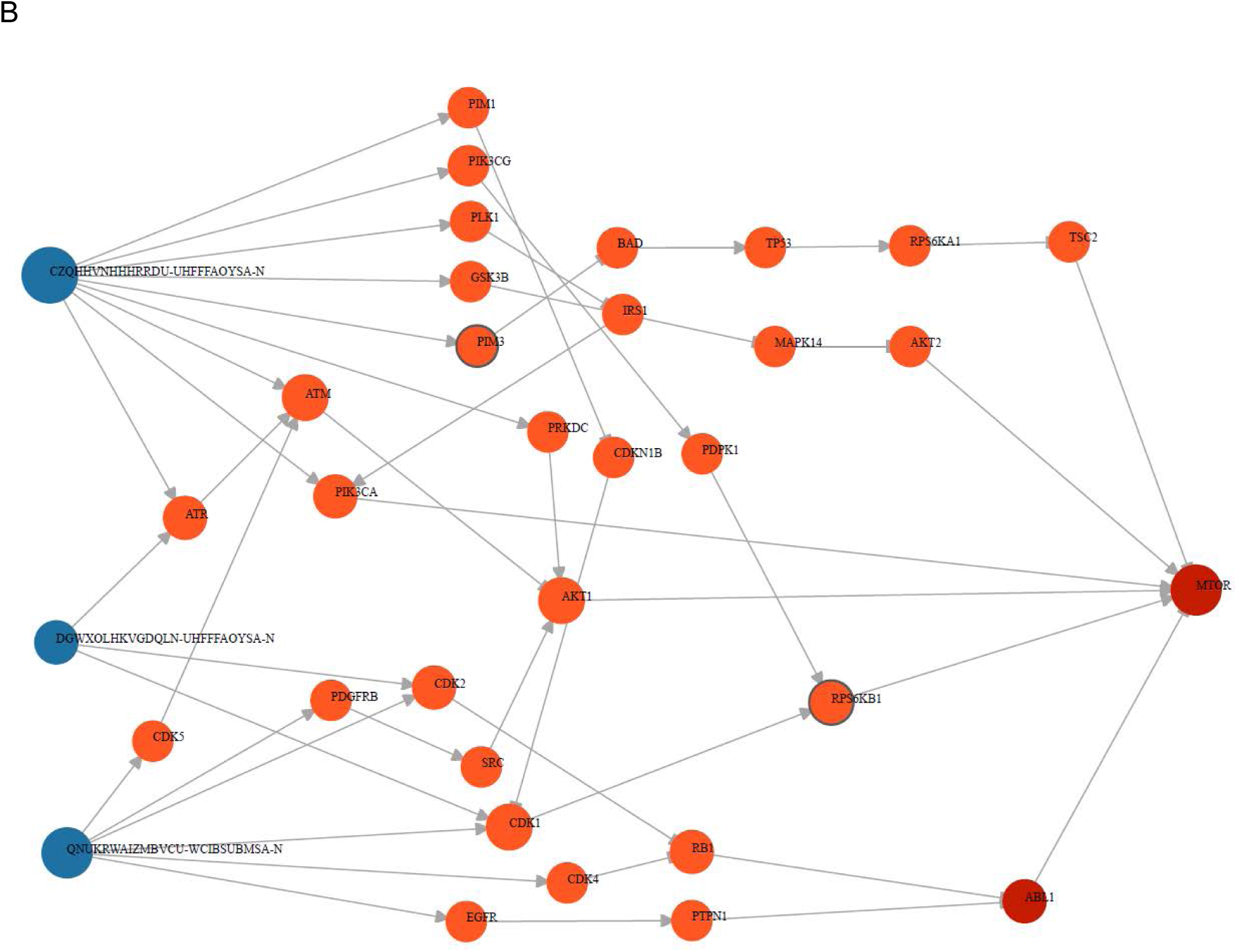
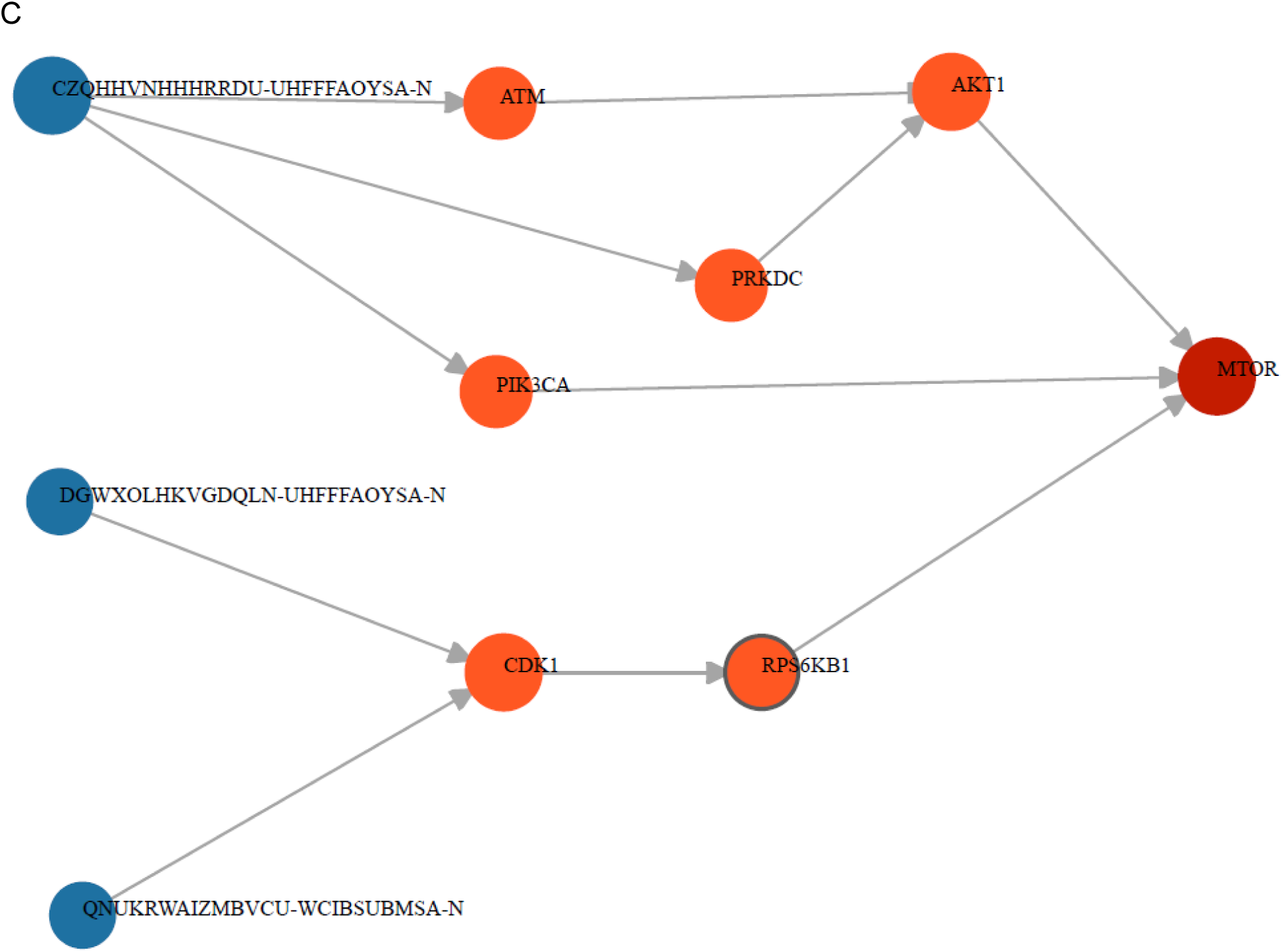
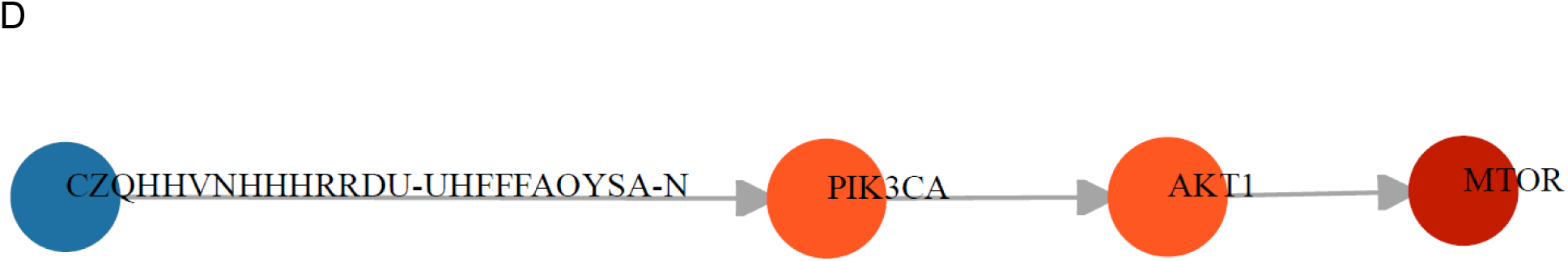
Second use-case. A. Shown is a Network generated by using the 8 active compounds (PubChem AID: 2660) as start nodes and mTOR as end node. Values of parameters k=5, p=3, c=0.01 were equal to their default values as they were not adjusted in the control panel. B. Showcasing weak interactions: two parameters were adjusted: k=5 and p=5. C. The value of k was set to 2 allowing for a maximal length of 2 of shortest paths (k=2, p=5, c=0.01). Other parameters were not adjusted. D. Generating subnetwork comprised of interactions of higher confidence based on curation. (k=10, p=5, c=0.73)

It is possible to restrict the network to only strong DTIs, which we can achieve by increasing the value of *p*, e.g. *p=5* (or 10μM) which sets a typical upper limit for active compounds. Consequently, one of the compounds (PubChem CID: 237) and paths induced by only that compound are eliminated from the subnetwork. Sometimes, it may be useful to limit the length of the shortest paths in order to eliminate distant (less certain) downstream effects. Accordingly, setting the value of *k* to *2* results in a smaller subnetwork. In this graph, the two compounds are more likely to perturb the action of the associated proteins.

Finally, the integration of an annotated and curated PPI database into the knowledge-base of SmartGraph (e.g., SIGNOR) allows the user to depict only PPI subnetworks with higher confidence. From a subnetwork generated with permissive parameter settings (*k=10*, *p=5*), we identified the maximal value of *c* that resulted in a subnetwork: *c=0.73*. Increasing the value of *c* further eliminates the “weakest link” between AKT1 and mTOR, consequently destroying the path between the only remaining compound and mTOR. This use-case might be preferable for someone who prioritizes curated data in MoA analysis. SmartGraph additionally provides the literature source (extracted from SIGNOR) of individual PPIs in the “Link View” of the information panel upon hovering the mouse over, or clicking on target-target edges.

In summary, the above scenarios allow the generation of hypotheses regarding the potential MoA of the active molecules. In these workflows, parameters *k*, *p* and *c* provide great versatility in prioritizing different aspects of the exploration process. Furthermore, the generated subnetwork can assist in analyzing how small molecules might perturb other targets in the network besides the end node(s).

### Bioactivity Prediction via Potent Patterns

The final aspect of SmartGraph is its integrated predictive functionality. As described in “Bioactivity Prediction” the investigator can predict potentially novel compounds for a target. In the frame of our example, bioactivity predictions can be made for mTOR as follows. By left-clicking on the mTOR node, then “Get predictions” in the revealed menu, compounds predicted to be associated with mTOR are shown. These compounds are clustered around potent patterns of mTOR that provided the basis for the prediction (see: *Figure 5a*). The overlap ratio between compounds and patterns can be obtained using “Link View” when mouse-hovering over or clicking on the respective compound-pattern edges. Generally, the higher the overlap ratio the more likely the predicted compound will be confirmed as a perturbagen of the target at hand. If the predicted compound is an approved drug, such a prediction could serve as the basis of a drug-repurposing study. When the number of predictions is large, one can perform a similarity-network based clustering of the predicted compounds to select a set of diverse compounds for further studies [41], [42], [43].

**Figure 5.**
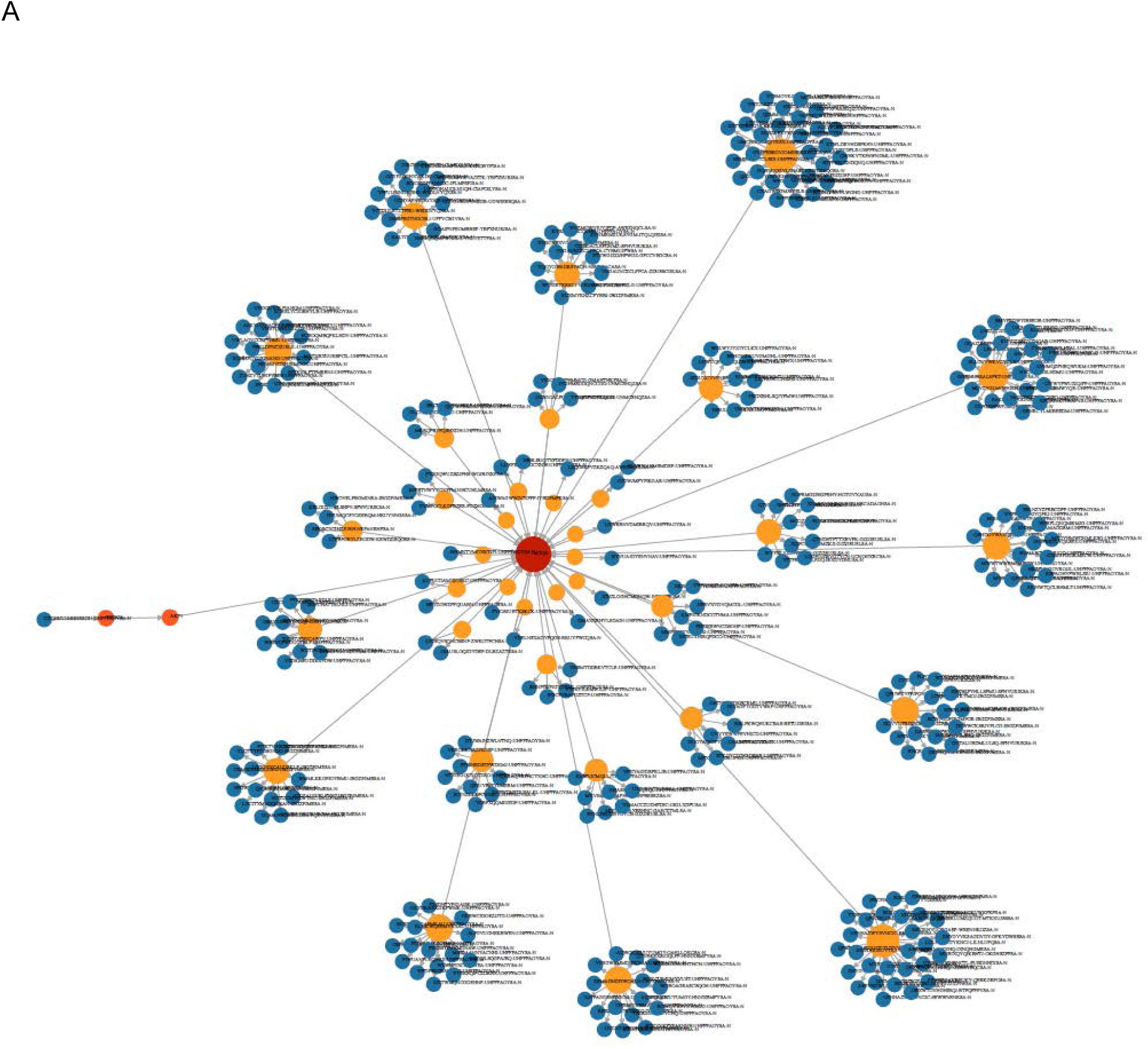
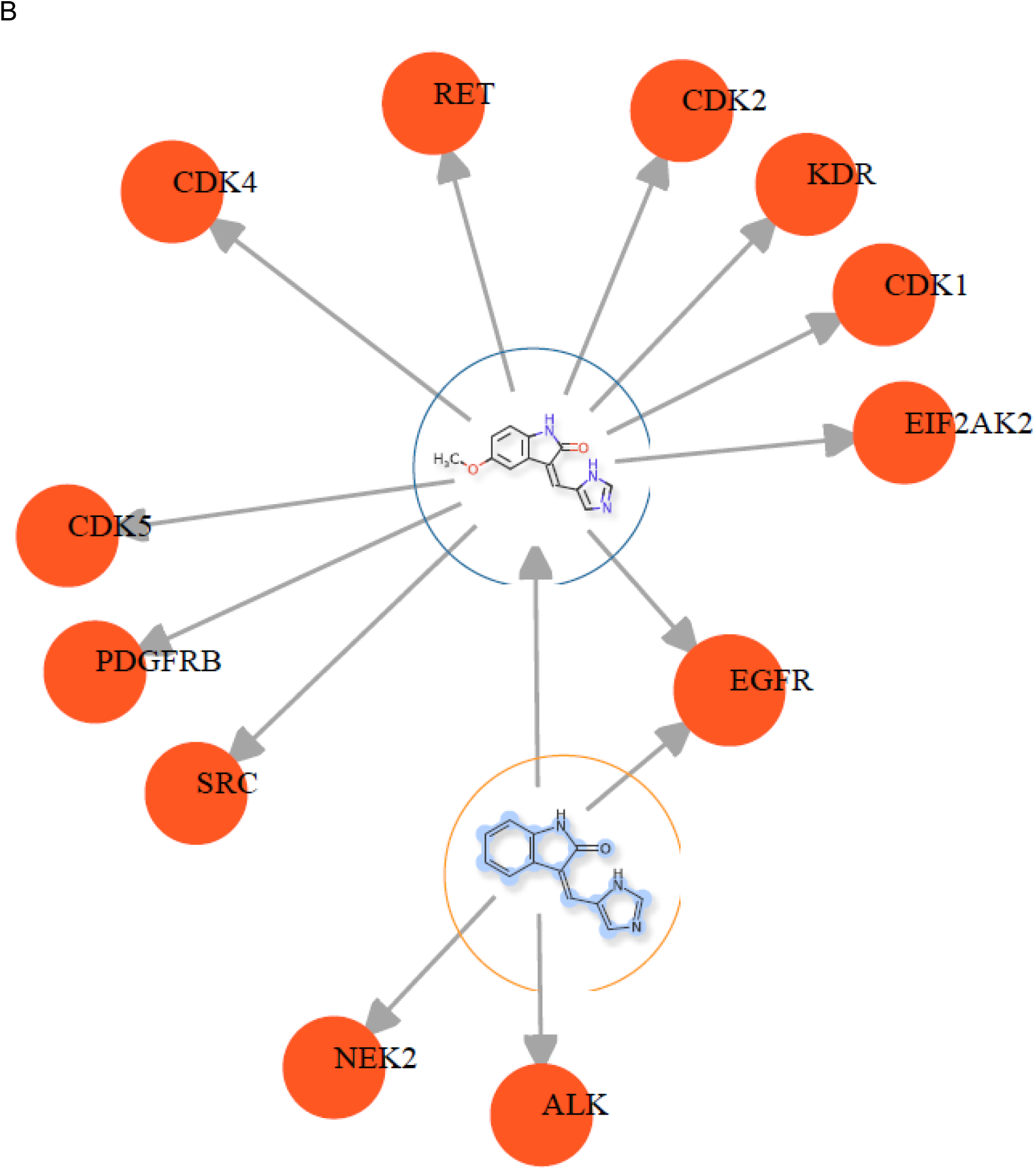
Third use-case. A. Result of prediction of active compounds for mTOR using the “Get Prediction” function of SmartGraph. Yellow nodes: patterns (BM-scaffolds), blue nodes: compounds, orange nodes: protein targets. This subnetwork is derived from the subnetwork generated by the last use-case of “Automated Hypothesis Generation for Mechanism-of-Action” section. B. Predicting polypharmacology for compound Compound A (NS-InChI key: QNUKRWAIZMBVCU, PubChem CID: 5289419). Scaffold overlap ratio: 0.89. Predicted targets: ALK (p68 kinase, P19525) and NEK2 (NimA-like protein kinase 1, P07949). Compound A was already tested in EGFR because it is connected to it.

Off-target interactions can be predicted in a similar fashion, i.e. taking advantage of the concept of potent patterns. In this use-case we use one of the active compounds from the above set of eight actives as a start node. Referred-to as Compound A (PubChem CID: 5289419, NS-InChI key: QNUKRWAIZMBVCU), the expansion results in 11 neighbor nodes (10 targets, one pattern) for Compound A (see: *Figure 5b*). Expanding the protein nodes of the pattern node reveals three targets (EGFR, NEK2, ALK) for which this is a potent pattern. Compound A was already tested on EGFR, but not on the other two targets. Thus, we hypothesize that Compound A is a modulator of NEK2 and ALK. The overlap ratio between the Bemis-Murcko scaffold and Compound A is 0.89, a reasonable probability for compound A being active on these two targets.

## Conclusions

In this study we presented SmartGraph, an integrated and predictive web-based platform which supports complex cheminformatics and bioinformatics workflows in a simple-to-use manner. The knowledge-base of SmartGraph is derived from ChEMBL v24 and the SIGNOR PPI databases, and contains 420,526 unique DTIs involving 271,098 unique compounds and 2,018 unique targets. Utilizing state-of-the-art technologies for backend (Neo4j) and frontend (Angular, RxJS, D3) designs, SmartGraph features powerful analysis workflows and data visualization. This was showcased through several use-cases that cover hypothesis generation for elucidating MoA, drug repurposing and off-target-prediction.

The design of SmartGraph and the underlying Neo4j graph-database allows for the facile integration of additional layers of biomedical data, such as pharmacological action of drugs, non-small molecule drugs, disease information and target development categories [44] to name a few. We also intend to integrate more cheminformatics and network analysis features into the platform in the future. Further, the graphical interface of SmartGraph can be used in many scenarios when efficient visualization is required to display, organize and analyze large-scale data. Finally, we plan to create a stand-alone API based on the GUI of SmartGraph and release it as an open-source code base.

SmartGraph is available at http://smartgraph.ncats.io/ upon the acceptance of this manuscript.

## Author’s Contributions

Gergely Zahoránszky-Kőhalmi, PhD (GZK) conceived and implemented the SmartGraph platform, Timothy Sheils designed and implemented the GUI. Tudor I. Oprea, MD PhD (TIO) conceived the CARLSBAD (chemical patterns) project and provided access to the CARLSBAD and KEGG databases which served as a first SmartGraph prototype. All of the authors contributed to the manuscript, GZK wrote most of the text and TIO rewrote it.

## Acknowledgments

This research was supported in part by the Intramural research program of the NCATS, NIH. Further, this study has been supported by the Illuminating the Druggable Genome – Knowledge Management Center NIH-U54 grant (1U54CA189205-01, PI: Tudor I. Oprea, MD PhD). The development of the CARLSBAD database was supported by the CARLSBAD NIH-R21 grant (GM095952, PI: Tudor I. Oprea, MD PhD).

Regarding the CARLSBAD-based prototype of SmartGraph, the authors are thankful for Stephen L. Mathias, PhD for his help in compiling active and inactive interactions from the CARLSBAD database, and for Jeremy J. Yang and Oleg Ursu, PhD for generating the chemical patterns for compounds of CARLSBAD database. Further, the authors are thankful to Ke Wang and Giovanni Torres, developers who helped to set up the server environment and infrastructure for SmartGraph.

## References

[1] A. L. Hopkins, “Network pharmacology.,” Nat. Biotechnol., vol. 25, no. 10, pp. 1110–1, Oct. 2007.

[2] T. I. Oprea et al., “Associating Drugs, Targets and Clinical Outcomes into an Integrated Network Affords a New Platform for Computer-Aided Drug Repurposing,” Mol. Inform., vol. 30, no. 2–3, pp. 100–111, Mar. 2011.

[3] T. I. Oprea and A. Tropsha, “Target, chemical and bioactivity databases – integration is key,” Drug Discov. Today Technol., vol. 3, no. 4, pp. 357–365, 2006.

[4] P. Shannon et al., “Cytoscape: a software environment for integrated models of biomolecular interaction networks.,” Genome Res., vol. 13, no. 11, pp. 2498–504, Nov. 2003.

[5] O. Taboureau et al., “ChemProt: a disease chemical biology database,” Nucleic Acids Res., vol. 39, no. suppl_1, pp. D367–D372, Oct. 2010.

[6] A. Roth et al., “STRING v10: protein–protein interaction networks, integrated over the tree of life,” Nucleic Acids Res., vol. 43, no. D1, pp. D447–D452, 2014.

[7] M. Kuhn, C. von Mering, M. Campillos, L. J. Jensen, and P. Bork, “STITCH: interaction networks of chemicals and proteins,” Nucleic Acids Res., vol. 36, no. Database issue, pp. D684–D688, Jan. 2008.

[8] R. J. Allaway, S. La Rosa, J. Guinney, and S. J. C. Gosline, “Probing the chemical-biological relationship space with the Drug Target Explorer,” J. Cheminform., vol. 10, no. 1, p. 41, Aug. 2018.

[9] L. Liang et al., “Bioactivity-explorer: a web application for interactive visualization and exploration of bioactivity data,” J. Cheminform., vol. 11, no. 1, p. 47, Dec. 2019.

[10] A. Venkat, PathInsight: A Novel Tool for Modeling Biomolecular Pathways. University of California, San Diego, 2017.

[11] D. P. Gil, J. N. Law, and T. M. Murali, “The PathLinker app: Connect the dots in protein interaction networks,” F1000Research, vol. 6, p. 58, Jan. 2017.

[12] http://datascience.unm.edu/carlsbad-platform/carlsbadone/, “CarlsbadOne.” [Online]. Available: http://datascience.unm.edu/carlsbad-platform/carlsbadone/.

[13] “Neo4j.” [Online]. Available: https://neo4j.com/.

[14] “Bolt Protocol.” [Online]. Available: https://boltprotocol.org/.

[15] “RxJS - Reactive Extensions Library for JavaScript.” [Online]. Available: https://rxjs-dev.firebaseapp.com/.

[16] “D3.js.” [Online]. Available: https://d3js.org/.

[17] T. Dwyer, “Scalable, Versatile and Simple Constrained Graph Layout,” Comput. Graph. Forum, vol. 28, no. 3, pp. 991–998, Jun. 2009.

[18] “Google Maps.” [Online]. Available: https://www.google.com/maps.

[19] G. Zahoránszky-Kõhalmi, T. I. Oprea, C. G. Bologa, S. Mani, and O. Ursu, “Network Inference Driven Drug Discovery,” University of New Mexico School of Medicine, 2016.

[20] G. W. Bemis and M. A. Murcko, “The properties of known drugs. 1. Molecular frameworks.,” J. Med. Chem., vol. 39, no. 15, pp. 2887–93, Jul. 1996.

[21] J. W. Raymond, E. J. Gardiner, and P. Willett, “RASCAL: Calculation of Graph Similarity using Maximum Common Edge Subgraphs,” Comput. J., vol. 45, no. 6, pp. 631–644, Jan. 2002.

[22] J. M. Zaretzki, M. K. Matlock, and S. J. Swamidass, “Scaffold network generator: a tool for mining molecular structures,” Bioinformatics, vol. 29, no. 20, pp. 2655–2656, 2013.

[23] S. J. Wilkens, J. Janes, and A. I. Su, “HierS: hierarchical scaffold clustering using topological chemical graphs.,” J. Med. Chem., vol. 48, no. 9, pp. 3182–93, May 2005.

[24] Jeremy J Yang, “Google Code open source project, unm-biocomp-hscaf, Java library for HierS chemical scaffolds.”.

[25] A. Gaulton et al., “ChEMBL: a large-scale bioactivity database for drug discovery,” Nucleic Acids Res., vol. 40, no. Database issue, pp. D1100–D1107, Jan. 2012.

[26] A. P. Bento et al., “The ChEMBL bioactivity database: an update.,” Nucleic Acids Res., vol. 42, no. Database issue, pp. D1083–90, Jan. 2014.

[27] A. Calderone et al., “SIGNOR: a database of causal relationships between biological entities,” Nucleic Acids Res., vol. 44, no. D1, pp. D548–D554, 2015.

[28] M. R. Berthold et al., “{KNIME}: The {K}onstanz {I}nformation {M}iner,” in Studies in Classification, Data Analysis, and Knowledge Organization (GfKL 2007), 2007.

[29] “PostgreSQL.” [Online]. Available: http://www.postgresql.org.

[30] “PostgreSQL JDBC Driver.” [Online]. Available: https://jdbc.postgresql.org/.

[31] “Neo4j JDBC Driver.” [Online]. Available: https://github.com/neo4j-contrib/neo4j-jdbc.

[32] S. Heller, A. McNaught, S. Stein, D. Tchekhovskoi, and I. Pletnev, “InChI - the worldwide chemical structure identifier standard,” J. Cheminform., vol. 5, no. 1, p. 7, Jan. 2013.

[33] T. U. Consortium, “UniProt: the universal protein knowledgebase,” Nucleic Acids Res., vol. 45, no. D1, pp. D158–D169, 2016.

[34] “Java - UUID.” [Online]. Available: https://docs.oracle.com/javase/1.5.0/docs/api/java/util/UUID.html.

[35] “Angular.” [Online]. Available: https://angular.io/.

[36] D.-T. Nguyen et al., “Pharos: Collating protein information to shed light on the druggable genome,” Nucleic Acids Res., vol. 45, no. D1, pp. D995–D1002, Nov. 2016.

[37] B. E. E., Y. Wang, P. A. Thiessen, and S. H. Bryant, “PubChem: Integrated Platform of Small Molecules and Biological Activities,” Annu. REPORTS Comput. Chem. VOL 4, vol. 4, pp. 217–241, 2010.

[38] P. M. L. Program, “Program, PubChem Molecular Libraries.”

[39] J. Inglese et al., “Quantitative high-throughput screening: A titration-based approach that efficiently identifies biological activities in large chemical libraries,” Proc. Natl. Acad. Sci., vol. 103, no. 31, pp. 11473–11478, 2006.

[40] “National Center for Biotechnology Information. PubChem Database. AID=2660.”

[41] L. A. Zahoránszky, G. Y. Katona, P. Hári, A. Málnási-Csizmadia, K. A. Zweig, and G. Zahoránszky-Köhalmi, “Breaking the hierarchy--a new cluster selection mechanism for hierarchical clustering methods.,” Algorithms Mol. Biol., vol. 4, no. 1, p. 12, Jan. 2009.

[42] G. Zahoránszky-Kőhalmi, C. G. Bologa, and T. I. Oprea, “Impact of similarity threshold on the topology of molecular similarity networks and clustering outcomes,” J. Cheminform., vol. 8, no. 1, p. 16, Dec. 2016.

[43] J. D. MacCuish and N. E. MacCuish, “Chemoinformatics applications of cluster analysis,” Wiley Interdiscip. Rev. Comput. Mol. Sci., vol. 4, no. 1, pp. 34–48, Jan. 2014.

[44] T. I. Oprea et al., “Unexplored therapeutic opportunities in the human genome,” Nat. Rev. Drug Discov., vol. 17, p. 317, Mar. 2018.

