## Supplementary Material for "SmartGraph: A Network Pharmacology Investigation Platform"

### Terminology

#### Nodes: Compound, Patterns and Targets

Compound nodes represent unique molecular structures extracted from substances of the ChEMBL database [1], [2] by removing salts and keeping only the largest component. They are attributed with an InChI [3] key and chemical structure in the SMILES [4], [5] format. The first section of the InChI key that does not encode stereochemistry is an additional attribute of compounds. This attribute is referred-to as non-stereo InChI key (NS-InChI key). Of note, the complete InChi key compared to the NS-InChi key, i.e. first-block of standard InChi key, in some cases might encode additional information beyond stereochemistry, e.g. different isotopes or presence of charge. However, in the context of the SmartGraph platform, using NS-InChi key is useful for identifying various stereoisomers of the same compound, considering the presence of ambiguous stereochemistry definitions of structures in large databases.

A chemical pattern, or pattern for short, is a useful concept in analyzing structure-activity relations as they represent common chemical features among compounds. In the current release of SmartGraph only Bemis-Murcko scaffolds (BM-Scaffolds) [6] were used as pattern, hence their pattern type attribute is ‘scaffold’.

Protein targets nodes are attributed with UniProt ID [7], protein name associated with the UniProt ID (part of ChEMBL 24.1 distribution)), gene symbol(s) (downloaded on: 11/05/2017) and synonym(s).

#### SmartGraph Relationships

*Tested – on.* The relation is defined between a compound C and a target node T and represents experimentally determined bioactivity, i.e. drug-target interaction (DTI). This relation is attributed with the aggregated $AC_{50}$/$IC_{50}$/$EC_{50}$ bioactivity values between the compound and target at hand, in the unit of $\mu$M. If multiple values are reported for the same C – T relation, then these values are aggregated as the median of such values. The edge representing this relation is directed: the start and end-nodes is C and T, respectively.

An important distinction was made between compounds in the context of this relation. That is, an activity threshold was defined to identify so-called *potent compounds* of a target. The activity cutoff attributes reflect the $80^{th}$ percentile of the potency values of all compounds tested on the target at hand. Of note, determining the activity cutoff involved all interactions regardless of the type of the bioactivity, e.g. if it is an $AC_{50}$, $IC_{50}$ or $EC_{50}$ value. The unit of the activity cutoff is -logM. If the number of interactions a target is associated with is <=5 than the activity cutoff was set to 7.

Of note, an additional attribute, e.g. type of activity, is also associated with DTI relations. However, currently SmartGraph includes both inhibitory and stimulatory relations, hence this attribute is set uniformly to: “activity”. In case other type of activity types are incorporated into the knowledge base this attribute can accommodate other type of activation data, e.g. inhibition, activation, dissociation constant.

*Regulates.* The relation is defined between two target nodes T1 and T2. The edge representing the relation is directed: the edge defined by start-node T1 and end-node T2 represents the regulatory relation where T1 regulates T2. PPI relations were extracted from the SIGNOR database [8].

*Pattern – of.* The relation is defined between a compound C and pattern P. In the current release of SmartGraph P is a Bemis-Murcko scaffold [6] of C. The edge representing the relation is directed: the start-node is P and the end-node is C. The *overlap ratio* is an attribute of the edge that gives the ratio between the number of heavy atoms in the P and C. The relation is attributed by the~~,~~ the overlap between the scaffold and compound expressed as the fraction of overlapping heavy atoms.

*Potent pattern – of.* Potent compounds of a target T are collected. Given a compound C in this collection, an associated pattern P of C is the potent pattern of T. Of note, it is possible that inactive compounds of the target at hand also contain this pattern. While this is a known phenomenon in the case of structure-activity relationship (SAR) series, SmartGraph utilizes a permissive strategy to maximize the likelihood of identifying active (potent) chemotypes.

DTI relations extracted from ChEMBL database are attributed by the internal identifier of the involved compound and target nodes, activity is expressed in unit of $\mu M$.

The direction of PPI and DTI relations is taken into account when analyzing the network as contrast to the direction of “pattern-of” and “potent-pattern-of” relations.

### Bioactivity Data Aggregation

Substances of ChEMBL database were converted to compounds by keeping only the largest component (CDK KNIME node [9], [10]). Next, InChI-keys and Morgan fingerprint [11], [12] of radius 3 and length of 2048 bits were generated for those compounds (RDKit KNIME node [13], [14]). Compounds associated with multiple fingerprints were removed from the data set (157 such compounds were found). The ChEMBL IDs of targets were resolved to UniProd IDs, leading to many-to-many relations. Accordingly, original DTIs (compound – target ChEMBL ID) were expanded to all possible compound-UniprotID tuples. In creating the knowledge base, only targets annotated as “SINGLE TARGET” according to the “chembl2uniprot” file, which is a part of the ChEMBL distribution, were used. Unique interactions and associated potency values were obtained by aggregating bioactivities according to their median values. In this process, we used the InChI keys and UniProt IDs to identify aggregate bioactivity data. Of note, only those bioactivities were aggregated that were associated with only one type of activity: $AC_{50}$, $IC_{50}$ or $EC_{50}$. Otherwise, the respective DTIs were ignored in deriving the knowledge base of SmartGraph.

BM-Scaffolds of compounds were detected in KNIME with the help of RDKit node. Scaffolds were deduplicated using ‘Group By’ KNIME Node and using the scaffold structure as key. InChI keys for scaffolds were generated by RDKit KNIME node.

| PubChem CID | InChi key | NS-InChi key | SMILES |
|---|---|---|---|
| 126280 | SVFXPTLYMIXFRX-XJKSGUPXSA-N | SVFXPTLYMIXFRX | CN[C@@H]1C[C@H](C2=CC=CC=C12)C3=CC(=C(C=C3)Cl)Cl |
| 2544 | NMTNUQBORQILRK-UHFFFAOYSA-N | NMTNUQBORQILRK | CC1(C2CCC(C1(C)C(=O)O)O2)C(=O)O |
| 398148 | DGWXOLHKVGDQLN-UHFFFAOYSA-N | DGWXOLHKVGDQLN | C1CCC(CC1)COC2=NC(=NC(=C2N=O)N)N |

| | | | |
|---|---|---|---|
| 237 | GPKJTRJOBQGKQK-UHFFFAOYSA-N | GPKJTRJOBQGKQK | CCN(CC)CCCC(C)NC1=C2C=C(C=CC2=NC3=C1C=CC(=C3)Cl)OC |
| 5289419 | QNUKRWAIZMBVCU-WCIBSUBMSA-N | QNUKRWAIZMBVCU | COC1=CC\2=C(C=C1)NC(=O)/C2=C\C3=CN=CN3 |
| 11957499 | HIYAVKIYRIFSCZ-CYEMHPAKSA-N | HIYAVKIYRIFSCZ | C[C@@H]1CC[C@]2([C@@H](C[C@H]([C@H](O2)[C@H](C)C(=O)C3=CC=CN3)C)C)O[C@@H]1CC4=NC5=C(O4)C=CC(=C5C(=O)O)NC |
| 11957469 | YARPKTMEHCQHGA-JPXKWFFHSA-N | YARPKTMEHCQHGA | C[C@H]1CCC/C=C/C2C[C@@H](C[C@]2(C/C=C/C(=O)O1)O)O |
| 3973 | CZQHHVNHHHRRDU-UHFFFAOYSA-N | CZQHHVNHHHRRDU | C1COCCN1C2=CC(=O)C3=C(O2)C(=CC=C3)C4=CC=CC=C4 |

**Table S1. Compounds involved in the use-cases.**

### References


[1] A. Gaulton *et al.*, “ChEMBL: a large-scale bioactivity database for drug discovery,” *Nucleic Acids Res.*, vol. 40, no. Database issue, pp. D1100–D1107, Jan. 2012.

[2] A. P. Bento *et al.*, “The ChEMBL bioactivity database: an update.,” *Nucleic Acids Res.*, vol. 42, no. Database issue, pp. D1083-90, Jan. 2014.

[3] S. Heller, A. McNaught, S. Stein, D. Tchekhovskoi, and I. Pletnev, “InChI - the worldwide chemical structure identifier standard,” *J. Cheminform.*, vol. 5, no. 1, p. 7, Jan. 2013.

[4] D. Weininger, “SMILES, a chemical language and information system. 1. Introduction to methodology and encoding rules,” *J. Chem. Inf. Model.*, vol. 28, no. 1, pp. 31–36, Feb. 1988.

[5] D. Weininger, A. Weininger, and J. L. Weininger, “SMILES. 2. Algorithm for generation of unique SMILES notation,” *J. Chem. Inf. Model.*, vol. 29, no. 2, pp. 97–101, May 1989.

[6] G. W. Bemis and M. A. Murcko, “The properties of known drugs. 1. Molecular

frameworks.," *J. Med. Chem.*, vol. 39, no. 15, pp. 2887–93, Jul. 1996.

[7] T. U. Consortium, "UniProt: the universal protein knowledgebase," *Nucleic Acids Res.*, vol. 45, no. D1, pp. D158–D169, 2016.

[8] A. Calderone *et al.*, "SIGNOR: a database of causal relationships between biological entities," *Nucleic Acids Res.*, vol. 44, no. D1, pp. D548–D554, 2015.

[9] C. Steinbeck, Y. Han, S. Kuhn, O. Horlacher, E. Luttmann, and E. Willighagen, "The Chemistry Development Kit (CDK): an open-source Java library for Chemo- and Bioinformatics," *J. Chem. Inf. Comput. Sci.*, vol. 43, no. 2, pp. 493–500, Mar. 2003.

[10] E. L. Willighagen *et al.*, "The Chemistry Development Kit (CDK) v2.0: atom typing, depiction, molecular formulas, and substructure searching," *J. Cheminform.*, vol. 9, no. 1, p. 33, Jun. 2017.

[11] H. L. Morgan, "The Generation of a Unique Machine Description for Chemical Structures-A Technique Developed at Chemical Abstracts Service.," *J. Chem. Doc.*, vol. 5, no. 2, pp. 107–113, May 1965.

[12] D. Rogers and M. Hahn, "Extended-connectivity fingerprints.," *J. Chem. Inf. Model.*, vol. 50, no. 5, pp. 742–54, May 2010.

[13] Greg Landrum, "RDKit: Open-source cheminformatics." [Online]. Available: http://www.rdkit.org/. [Accessed: 24-Feb-2018].

[14] "RDKit Nodes for KNIME." [Online]. Available: https://www.knime.com/nodeguide/community/rdkit.
